# Adaptation of *Mimulus cardinalis* to drought through changes in stomatal conductance

**DOI:** 10.64898/2026.11.18.763328

**Authors:** Dachuan Wang, Lucas J. Albano, Charlie Blanchard, Lauryn Cabral, Emily Cook, Christina Cranwell, Lily Ruth Danz, Rosalinda Diaz, Xitlaly Gomez-Vega, Aren Gonzales, Tony Gurnoe, Katelin Kutella, Mariam Moazed, Macy Patel, Julia Prange, Niveditha Ramadoss, Ashley Regan, Aster Riley, Melissa Rivas Hernandez, Jocelyn Rojas, Caitlyn Shank, Marissa Strebler, Gabriel Tejeda, Genevieve Triplett, Aditi Verma, Lindsay Villano, Jordan Waits, Olivia Wilborn-Pilotte, Jason P. Sexton, Jeffrey Diez, Lluvia Flores-Rentería, Seema Nayan Sheth, Christopher D. Muir

## Abstract

**Premise of the study:** Many areas around the globe are experiencing warmer temperatures, more extreme drought, and wildfires because of climate change. Therefore, plant populations may experience strong natural selection on traits associated with adaptation to rapid change. One trait likely under selection is stomatal conductance, which determines the rate at which water vapor and carbon dioxide diffuse through the stomata and is sensitive to environmental factors such as drought, CO_2_, and vapor pressure deficit. The purpose of this study is to understand how plants respond both ecologically and evolutionarily to an extreme multiyear drought.

**Methods:** Using six populations of a perennial herb (*Mimulus cardinalis*), we measured the evolution of operational stomatal conductance in pre-drought ancestral and post-drought descendant cohorts planted in three common gardens across the species’ range. This design allowed us to test (1) whether stomatal conductance evolved in response to drought and (2) whether populations with greater genetic variance evolved more.

**Results:** Patterns consistent with drought escape were more common than patterns consistent with avoidance (6 versus 4 of 18 population-by-garden combinations), but evidence for evolutionary change in stomatal conductance was limited overall, and the extent of change was not correlated with genetic variance.

**Conclusions:** These findings offer a framework for future research on stomatal traits and insight into the rate at which plants respond to environmental change.

## INTRODUCTION

Climate change increases drought severity and aridity in many regions, creating strong and rapid selection on plant populations (Franks et al., 2007). Plants are the primary producers on Earth, generating organic matter through photosynthesis, and many traits, such as growth, are sensitive to environmental change (Driesen et al., 2020). For example, during the Quaternary, atmospheric CO_2_ remained low (170–210 ppm) during glacial maxima over the past 3.6 million years (Berends et al., 2021), but human activities have rapidly raised global mean CO_2_ to about 430 ppm (Tripati et al., 2009; Becklin et al., 2017; NOAA GML CCGG Group, 2019). Elevated CO_2_ contributes to warming, greater drought severity, and drier atmospheres (Ficklin and Novick, 2017). Temperature and water stress could affect plants both directly and indirectly (Leakey et al., 2009; Becklin et al., 2017). Across a species’ range, populations occupy habitats that differ in environmental pressures, shaping local adaptation, standing genetic variation, and plasticity. Populations from historically variable climates may exhibit greater adaptive plasticity (Levins, 1968; Rehfeldt et al., 1999; Sexton et al., 2017), which can maintain fitness across environments but weaken selection (Ghalambor et al., 2007). In contrast, populations in stable climates may have reduced plasticity and stronger local specialization (Levins, 1968; Kawecki and Ebert, 2004; Sexton et al., 2017), and they may evolve faster under consistent selection if standing variation is available (Ghalambor et al., 2007; Salamin et al., 2010). Overall, changing environmental conditions can act as agents of natural selection, leading to rapid adaptive change in plant populations (Franks et al., 2007).

Stomata are microscopic pores, usually on the leaf surface, each formed by two guard cells. They are the main pathway for gas exchange between plants and the atmosphere. Stomatal conductance (*g*_s_) quantifies the conductance of stomata to water vapor or CO_2_ diffusion, integrating stomatal aperture, density, and size. To assess stomatal responses to drought, we measured stomatal conductance to water vapor (*g*_sw_, mol m^−2^ s^−1^) under ambient conditions, which is often referred to as “operational” stomatal conductance (Murray et al., 2020). Stomata generally do not open maximally because of constraints such as insufficient soil water, risk of xylem cavitation, or stress-induced non-stomatal limitations on photosynthesis (Givnish and Vermeij, 1976; Flexas et al., 2012; Drake et al., 2017). Plants regulate stomatal conductance through both plastic and evolutionary responses. Multiple mechanisms can affect stomatal opening and closing, and these responses can occur over short or long timescales (Hetherington and Woodward, 2003). When water supply is insufficient, plants close their stomata, causing a short-term decrease in *g*_s_. Over the long term, evolutionary responses may reduce stomatal density, stomatal size, and the anatomical maximum stomatal conductance across several generations (Cowan and Farquhar, 1977; Woodward, 1987; Royer, 2001; Morgan et al., 2004; Buckley and Mott, 2013; Wang et al., 2020). Other factors, such as high vapor pressure deficit (VPD), increase rates of water loss and cause more drought stress (Running, 1976; Hetherington and Woodward, 2003). Plants respond not only physiologically through changes in aperture; stomatal density and size may also change plastically or evolutionarily. Environmental signals and endogenous plant hormones control stomatal development (Qi and Torii, 2018).

When plants face drought stress, they respond with strategies such as drought escape and drought avoidance (Abobatta, 2019). Drought escape refers to the ability of plants to complete their life cycle before the onset of severe water deficit, typically through early flowering and seed set (Levitt, 1980). Drought avoidance involves maintaining high tissue water potential under water-limited conditions to help plants survive long, dry seasons. This is usually achieved through reduced photosynthesis and improved root water uptake (Levitt, 1980; Basu et al., 2016). These contrasting strategies involve different developmental traits and timing, including differences in stomatal traits (Kooyers, 2015). For example, if plants employ a drought escape strategy, we expect natural selection to favor higher *g*_sw_, particularly early in the growing season, to accelerate development. As the season becomes increasingly dry, *g*_sw_ may then decline in response to the effects of drought. In contrast, if plants employ a drought avoidance strategy, natural selection should favor lower *g*_sw_ early in the season, allowing them to conserve water and withstand the harsh effects of heat and drought later in the growing season. However, because drought-avoiding plants take longer to complete their life cycle, *g*_sw_ may increase later in the season, The black/grey dots indicate different seasons. as they must sustain photosynthesis and growth during the dry season (Kooyers, 2015; Anstett et al., 2021; Branch et al., 2024b). These predictions are illustrated in Fig. 1.

**Figure 1:**
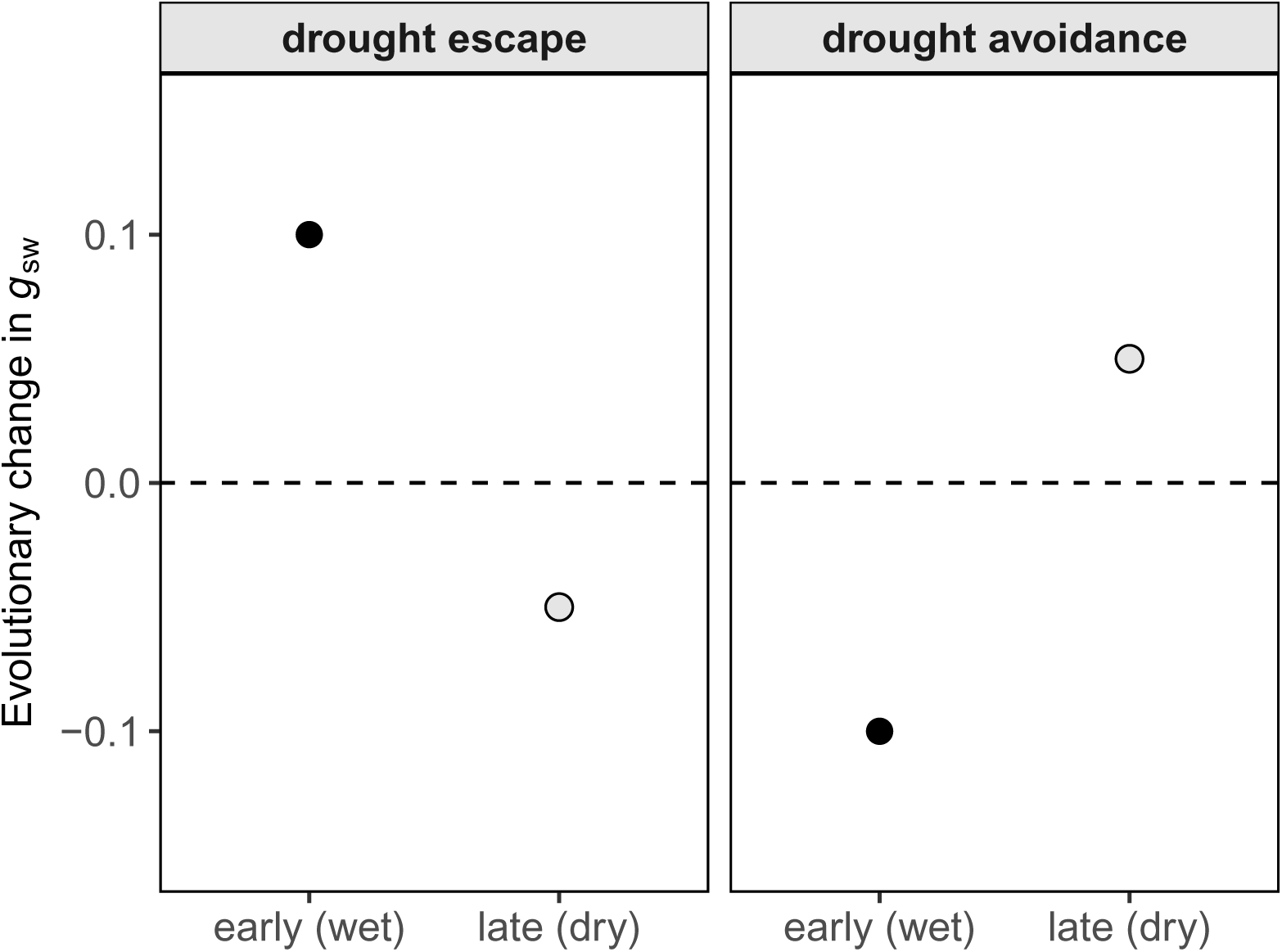
Simulated schematic diagram of stomatal conductance (*g*_sw_) under two different strategies.

Whether populations evolve toward either strategy depends not only on selection but also on the heritable variation available. Under the breeder’s equation, the response to selection is the product of narrow-sense heritability and the strength of selection (*R* = ℎ^2^*S*) (Lande and Arnold, 1983; Falconer and Mackay, 1996), so populations with greater heritability should evolve faster when selection is similar. Genetic variation may also differ across the species’ range; for example, genetic variation in phenology in *M. cardinalis* is greater at the trailing edge (Sheth and Angert, 2016).

*Mimulus* (*Erythranthe*) *cardinalis* is a perennial herb found across a broad climatic and latitudinal gradient from central Oregon to northern Baja California (Fraga, 2018; Lowry et al., 2019). It has long been used to study adaptation to environmental stress and the geographic limits of its range (Hiesey et al., 1971; Schemske and Bradshaw, 1999; Angert and Schemske, 2005; Angert, 2006a, b; Angert et al., 2018), and leaf physiological and life-history traits show substantial heritable variation (Nelson et al., 2021; Muir et al., 2022). Because individuals can flower and germinate every year (Sheth and Angert, 2018; Anstett et al., 2026), evolutionary responses may be detectable over relatively short timescales. During the early 2010s, western North America experienced severe drought (Diffenbaugh et al., 2015; Robeson, 2015; Wang et al., 2016) that varied geographically across the range of *M. cardinalis* and caused widespread population declines, especially where winters were driest (Sheth and Angert, 2018; Anstett et al., 2026). This created spatially heterogeneous selection and an opportunity to test for evolutionary responses with a resurrection approach, which compares ancestral and descendant genotypes derived from seeds collected before and after a period of environmental change (Franks et al., 2007, 2018). Previous resurrection studies in this species found that flowering time did not advance (Vtipil and Sheth, 2020) and that heritabilities of several escape and avoidance traits were low and variable among populations (Sheth et al., 2026), suggesting that constraints may limit rapid evolution. Resurrection studies of *M. cardinalis* in greenhouse conditions also measured stomatal conductance and found regional differences in the evolution of leaf traits following the same drought (Branch et al., 2024b), but evolutionary change in stomatal conductance has not been tested under field conditions.

To test these hypotheses, we asked the following research questions: (1) How does *g*_sw_ vary across early (wet) and late (dry) seasons, source populations, and the geographic range in three gardens of *M. cardinalis*? (2) Did *g*_sw_ evolve between 2010 and 2017 in the direction predicted by selection on drought escape or drought avoidance? (3) Was the magnitude of evolutionary change among populations and gardens correlated with heritability of *g*_sw_? This study examines both evolutionary and plastic responses of *M. cardinalis* to drought, revealing how *g*_sw_ varies across generations, gardens, and seasons.

## METHODS

### Study system

We used *Mimulus* (*Erythranthe*) *cardinalis* to investigate how stomatal conductance evolves across a species’ range in response to a major drought. We used a resurrection approach in common gardens (Fig. 2A) to estimate evolved genetic differences between ancestral and descendant cohorts, and we used the half-sib family structure of the experiment to estimate genetic variance in stomatal conductance in the ancestral cohort.

**Figure 2:**
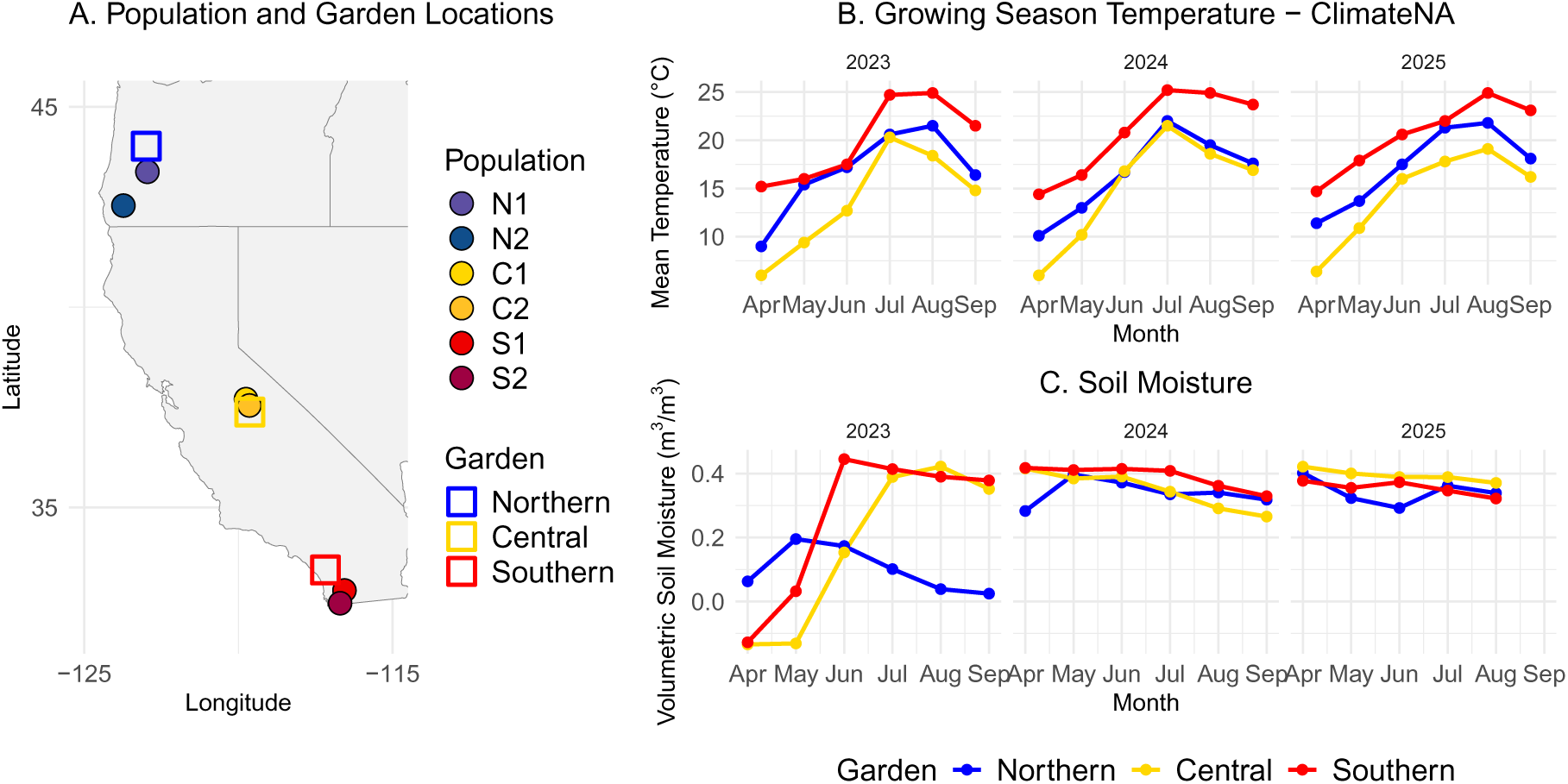
A) Map showing the known *Mimulus cardinalis* focal populations (filled circles) and common garden locations (squares). B) Mean monthly growing-season temperature (°C; April– September) for each common garden across three years (2023–2025), derived from ClimateNA climate data (v7.30; Wang et al., 2016; http://climatena.ca). C) Volumetric soil moisture (m^3^ m^−3^) dynamics for each garden across three years (2023–2025).

### Experimental design

We used a resurrection (Franks et al., 2018) and common garden experiment to compare plant traits among garden sites, source populations, and cohorts, following established approaches in this system (Sheth and Angert, 2016; Wooliver et al., 2020; Sheth et al., 2026). This experiment was previously described in Sheth et al. (2026); we summarize the key details here. We selected six source populations (filled circles in Fig. 2A): two leading-edge (northern), two central, and two trailing-edge (southern) populations of *M. cardinalis* (Sheth and Angert, 2016, 2018; Wooliver et al., 2020; Sheth et al., 2026). For each source population, we used seeds from $ individuals. Seeds were collected for each population in 2010 (ancestral cohort) and 2017 (descendant cohort). These field-collected seeds were grown in a greenhouse at North Carolina State University for one generation to control for maternal and storage effects (following Franks et al., 2007). Because population sizes differ across the species’ range, the number of parental individuals also varies among populations (Sheth et al., 2026).

We germinated pedigreed seeds for each population in greenhouses at the University of Oregon, the University of California, Merced, and San Diego State University in 2023. We kept all population-cohort combinations at similar sample sizes by using the same number of individuals in each group (source population × cohort). Based on the varying number of available full-sib families per group, we adjusted the number of replicates for each full-sib family. At 4 to 6 weeks after sowing, we transplanted 14,290 plants in total (an average of 21 sires and 101 dams per ancestral cohort of each population) into the gardens in a randomized block design. Within each garden, plants from each source population and cohort were randomly assigned to blocks, and their positions within blocks were also randomized.

We established common gardens in three sites within the range of *M. cardinalis*: near the northern range limit (Friends of Buford Park and Mt. Pisgah’s Native Plant Nursery; Eugene, OR), the latitudinal center (Sierra National Forest’s Batterson Work Center; Oakhurst, CA), and the southern range limit (San Diego State University’s (SDSU) Santa Margarita Ecological Reserve; Temecula, CA; Fig. 2).

### Stomatal conductance measurements

We measured operational stomatal conductance to water vapor (*g*_sw_) in three years (2023, 2024, and 2025) in each of two seasons (early and late). The early season occurred from early May to late June, when most individuals were in the active vegetative growth stage. The late season was in August, by which time plants had already passed peak flowering. See Table S1 for the detailed schedule. Measurements were taken using handheld porometers (LI-600, LI-COR Environmental, Lincoln, NE, USA). In all three gardens, *g*_sw_ was measured on a single leaf per plant.

We conducted measurements on clear days between 9:00 a.m. and 5:00 p.m. We waited for the dew to evaporate fully to avoid affecting porometer readings, and we began measurements once direct light intensity exceeded 1200 *μ*mol m^−2^ s^−1^. Under laboratory conditions, plants reached near-maximal *g*_sw_ at this light level (data not shown). We developed a standard protocol for leaf selection and documented any necessary adjustments. In the early season, measurements were conducted on the uppermost fully expanded, minimally shaded, healthy, undamaged, and dry leaf on the main stem, ensuring the leaf filled the instrument aperture. On some short plants, all leaves were close to the ground, so we measured the least shaded side. When damage was unavoidable because of herbivory, we selected the least-damaged leaf. In the late season, the same criteria were applied; however, for large or stressed individuals, priority was given to the healthiest available green leaves, relaxing the selection criteria if necessary but avoiding completely dry or yellow leaves. When multiple suitable leaves were available, preference was given to higher, uncovered leaves and to leaves on main branches.

### Garden climate and soil moisture

To describe conditions at the gardens (Fig. 2B, C), we obtained monthly mean temperature for the location of each garden from ClimateNA version 7.30 (Wang et al., 2016) for April to September of 2023 to 2025. We measured soil moisture at 10 cm soil depth in each garden with TOMST data loggers (Wild et al., 2019), one logger per block. We converted the loggers’ sensor counts to volumetric soil moisture (m^3^ m^−3^) using a quadratic calibration equation from the manufacturer and calculated monthly means for April to September of 2023 to 2025. September 2025 is omitted because only the northern garden has data for that month.

### Statistical analyses

All statistical analyses were performed in *R* version 4.6.0 (R Core Team, 2024). We used the **brms** package version 2.23.0 (Bürkner, 2017, 2018) to analyze the stomatal conductance data. This method employs Hamiltonian Monte Carlo (HMC) via *Stan* version 2.38.0 (Carpenter et al., 2017) and is implemented using the **cmdstanr** backend version 0.9.0 (Gabry et al., 2024). We used the modified Gelman-Rubin *R̂* statistic (Vehtari et al., 2021) to assess parameter convergence. For each garden-specific model, we ran four parallel HMC chains. For the southern and northern garden models, each chain consisted of 4,000 total iterations, with the first 2,000 iterations discarded as warm-up, leaving 2,000 sampling iterations per chain (thinning interval of 1). However, the central garden model did not converge under these settings. We used 8,000 total iterations per chain, with the first 4,000 iterations as warm-up and 4,000 sampling iterations per chain, with a thinning interval of 2 to reduce autocorrelation. Parameter estimates and credible intervals were summarized using the median and the 95% quantile intervals of the posterior distribution.

We analyzed differences in *g*_sw_ between ancestral and descendant cohorts of each source population by fitting a separate model to the data from each garden. *g*_sw_ (mol m^−2^ s^−1^) was modeled with a Gaussian distribution and identity link. The fixed effects were source population, cohort (ancestral or descendant), season (early or late), collection year (2023, 2024, or 2025), leaf vapor pressure deficit at the time of measurement (kPa), and time of day with a quadratic term to account for non-linear responses. Time of day was centered and scaled to improve numerical stability. We also included the two-way interactions population × cohort, population × season, population × year, cohort × season, and season × year, and the three-way interactions population × cohort × season and population × season × year. The random effects were sire and dam nested within sire, each with an intercept and effects of population, season, and their interaction (all uncorrelated), and block and date (intercepts). The residual variance was allowed to differ among populations, seasons, and their interaction. The models were fitted to 44,771 measurements (21,723 in the northern garden, 9,792 in the central garden, and 13,256 in the southern garden) from 13,481 plants with known population and cohort.

### Variation in g_sw_ among populations, cohorts, and seasons within gardens

To summarize variation in *g*_sw_ among populations, seasons, and years within and among gardens, we used each model to predict *g*_sw_ for plants of the ancestral cohort of every population in each season and year. Predictions were made at the mean VPD and time of day measured in that garden, season, and year, and with the random effects set to zero (population-level predictions). No prediction was possible for the early season of 2023 in the central garden because no measurements were made. We summarized predictions by their posterior mean and 50% and 95% quantile intervals (Fig. 3, Table S2). To compare populations, we calculated the difference in predicted *g*_sw_ between every pair of populations within each garden, season, and year, and between every pair of garden-year combinations within each population and season, for each posterior draw (Fig. S1, Fig. S2). A difference was considered significant when its 95% credible interval did not overlap zero. To compare southern (S1, S2) and northern (N1, N2) populations in the southern garden, we averaged the predicted *g*_sw_ of the two populations in each region for each posterior draw before taking the difference. We identified the garden with the highest mean *g*_sw_ in each season and year from the mean of the posterior mean predictions across populations. Other parameter estimates and uncertainty were summarized using posterior medians and 95% credible intervals.

**Figure 3:**
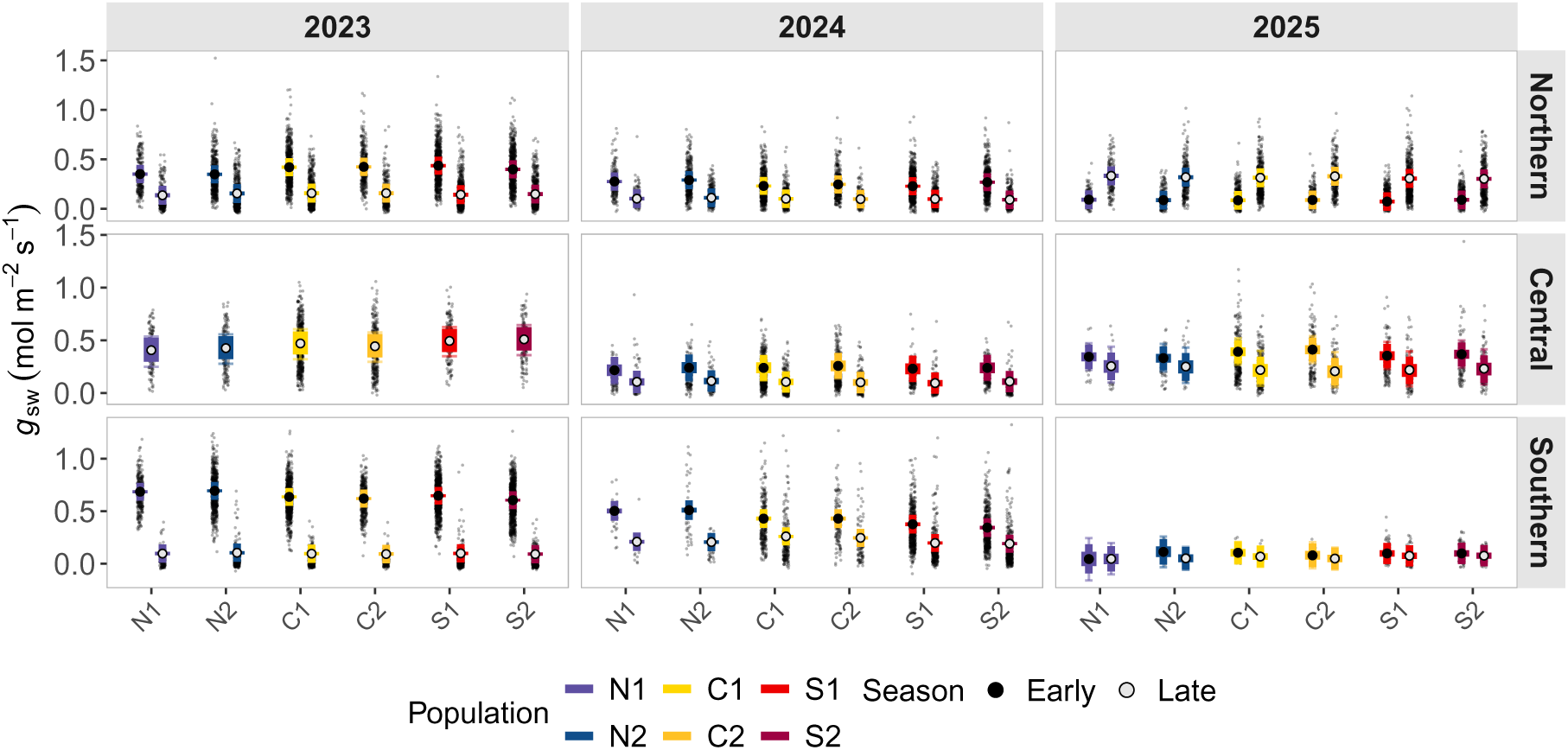
Posterior predictions of stomatal conductance (*g*_sw_, mol m^−2^ s^−1^) for early (lighter) and late (darker) seasons of each source population from the northern range edge (N1, N2), latitudinal range center (C1, C2), and southern range edge (S1, S2) in each common garden. White-filled points represent posterior means, thick lines indicate 50% credible intervals, and thin lines indicate 95% credible intervals. Raw data are overlaid as semi-transparent points in the background.

### Analyses of evolutionary change in stomatal conductance (g_sw_)

To determine whether changes in *g*_sw_ reflected escape or avoidance strategies, we compared responses across early and late season conditions. Evolutionary responses in *g*_sw_ were quantified as the posteriordifference between descendant and ancestral cohorts for all populations in each garden and season, predicted as above for 2024 (because the models have no cohort × year interaction, the difference is the same in every year):

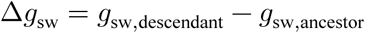

We classified evolutionary change as consistent with escape or avoidance by comparing the posterior means of Δ*g*_sw_ between the early and late seasons:

- Escape strategy: Δ*g*_sw,early_ > 0 and Δ*g*_sw,late_ < 0
- Avoidance strategy: Δ*g*_sw,early_ < 0 and Δ*g*_sw,late_ > 0

We did not include a cohort × collection year interaction in the model because mortality reduced the sample size in 2025, which in turn reduced the statistical power to detect differences between ancestral and descendant cohorts across collection years. Populations that did not follow either pattern were considered to show no clear adaptive response. Evolutionary change was considered significant when the 95% credible interval of the posterior contrast did not overlap zero. We summarized Δ*g*_sw_ by its posterior mean and 50%, 80%, and 95% quantile intervals (Fig. 4).

**Figure 4:**
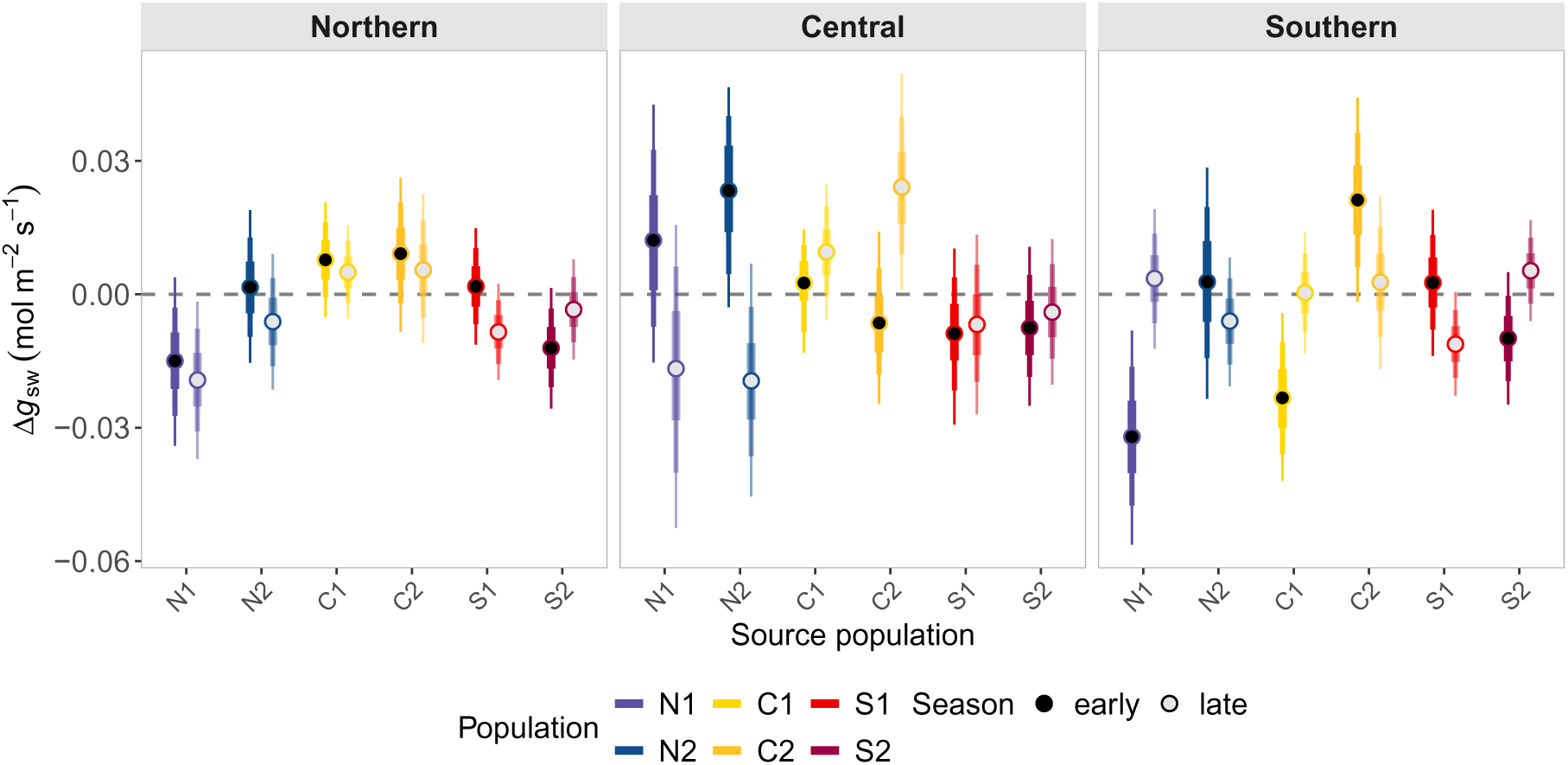
Δ*g*_sw_ (mol m^−2^ s^−1^; descendant minus ancestral cohort) for each population and garden. Solid circles show posterior means in the early season, open circles show posterior means in the late season, and thick, medium, and thin error bars show 50, 80, and 95% credible intervals.

#### Narrow-sense heritability

For each population and season, we estimated the additive genetic variance (*V*_A_) and the phenotypic variance (*V*_P_) of *g*_sw_ from the variance components of the model: the sire variance 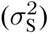, the variance among dams within sires 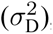, and the residual variance 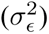 (Falconer and Mackay, 1996; Conner and Hartl, 2004). We use these estimates as the genetic variance of the ancestral cohort, the population at the start of the drought. In a nested half-sib design, 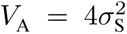 and 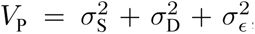, so narrow-sense heritability is 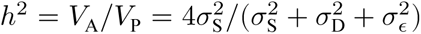. Because the sire and dam random effects of population, season, and their interaction were uncorrelated, 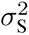 and 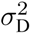 for a given population and season are the sums of the variances (squared standard deviations) of the terms that apply to that population and season. 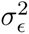 is the squared residual standard deviation for that population and season. We calculated ℎ^2^ for each posterior draw. We summarized ℎ^2^ by its posterior median and 50%, 80%, and 95% quantile intervals (Fig. 5). To compare seasons, we calculated the difference in ℎ^2^ between the late and early seasons for each posterior draw within each population and garden, and considered the difference significant when its 95% credible interval did not overlap zero. We estimated ℎ^2^ for all populations in the northern and southern gardens but only for C1 and C2 in the central garden (see below).

**Figure 5:**
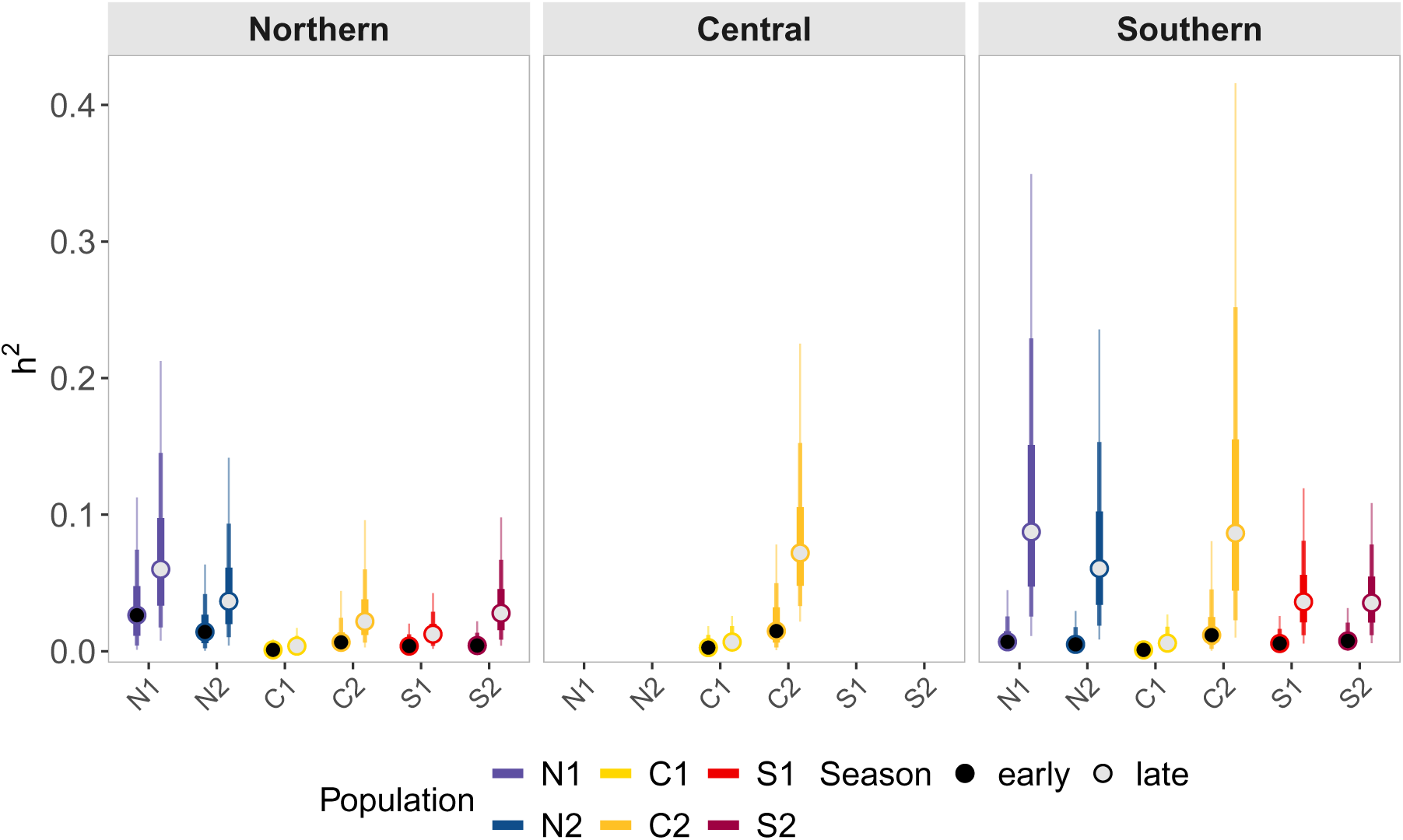
Narrow-sense heritability (ℎ^2^) of each population in each garden. Solid circles show posterior medians in the early season and open circles show posterior medians in the late season. The three nested intervals show 50%, 80%, and 95% credible intervals, from innermost to outermost.

#### Relationship between evolutionary change and heritability

To assess whether evolutionary change in Δ*g*_sw_ was associated with ℎ^2^, we quantified their relationship across populations using posterior samples. We paired the posterior draws of Δ*g*_sw_ and ℎ^2^ for each population-by-garden combination and performed linear regressions of Δ*g*_sw_ on ℎ^2^ separately for each season and each posterior draw, pooling all population-by-garden combinations. Due to space constraints in the central garden, we were unable to plant enough individuals to estimate ℎ^2^ of the southern and northern source populations. This left 14 population-by-garden combinations (data from the southern and northern populations in the central garden were excluded). This yields a posterior distribution of regression slopes for each season. We then summarized each slope distribution by its posterior mean, 95% credible interval, and the posterior probability that the slope exceeds zero, providing evidence for the direction and strength of the relationship between evolutionary change and heritability.

## RESULTS

### Variation in *g*_sw_ among populations, cohorts, and seasons within gardens

Stomatal conductance (*g*_sw_) differed strongly among gardens and seasons, with a consistent seasonal pattern of higher *g*_sw_ in the early than in the late season in most gardens and years (Fig. 3; Table S2). This pattern was clearest in the early season in the northern and southern gardens in 2023 and 2024, where early season *g*_sw_ was substantially higher than late season *g*_sw_. In the central garden, early season *g*_sw_ was also higher than late season in 2024 and 2025, but early season measurements were not available for 2023 because delayed transplanting, caused by late, heavy snowfall, left plants too small to measure in May and June. The seasonal pattern was different in 2025 in two gardens. In the northern garden, the pattern reversed, meaning that late season *g*_sw_ was higher than early season *g*_sw_ in all populations. In the southern garden, the early season values were not as elevated as in previous years. The central garden showed no such change in 2025 (Table S2 gives detailed values).

Beyond the differences between seasons and among gardens, *g*_sw_ also varied among populations and collection years. Comparing the same populations across gardens, mean *g*_sw_ was highest in the southern garden in the early season of 2023, the early season of 2024, and the late season of 2024, in the central garden in the late season of 2023 and the early season of 2025, and in the northern garden in the late season of 2025. Between the northern and central gardens, some populations did not differ clearly in this trait, whereas others differed significantly (Fig. 3; Table S2). The magnitude of these differences was also not consistent across seasons. Among populations within the same garden, a higher proportion of pairwise contrasts was significant in the early season (49.2%) than in the late season (23.7%) (Fig. S1). Across collection years, early season *g*_sw_ in the northern garden declined from 2023 to 2025 (Fig. 3). For the central garden, *g*_sw_ trended upward from 2024 to 2025, but the trend was not significant. In contrast, early season *g*_sw_ decreased over the three years in all populations in the southern garden. For late season *g*_sw_, the northern garden showed higher values in 2025 than in the previous two years. In the central garden, late season *g*_sw_ declined from 2023 to 2024 in all populations but did not change significantly between 2024 and 2025. In the southern garden, late season *g*_sw_ was higher in 2024 than in 2023 and 2025, which did not differ significantly from each other.

### Evolutionary change in *g*_sw_ from ancestral to descendant cohorts across populations and gardens

We found that responses from ancestral to descendant cohorts were diverse across populations and gardens (Fig. 4). Of the 18 population-by-garden combinations, 6 showed a pattern consistent with a shift toward an escape strategy (increasing early season and decreasing late season *g*_sw_): N2 in the northern garden, S1 in the northern garden, N1 in the central garden, N2 in the central garden, N2 in the southern garden, and S1 in the southern garden. For example, population N2 showed this pattern in all three gardens. A further 4 combinations more closely resembled the evolution of an avoidance strategy (decreasing early season and increasing late season *g*_sw_): C2 in the central garden, N1 in the southern garden, C1 in the southern garden, and S2 in the southern garden. However, for some populations, the response trends were less clear. In the northern garden, populations C1 and C2 showed increasing *g*_sw_ from ancestral to descendant cohorts in both early and late seasons, whereas N1 and S2 decreased in both seasons. In the southern garden, most populations showed minimal changes between cohorts. Only 4 of the 36 changes had 95% credible intervals that excluded zero: N1 in the northern garden in the late season (decrease), C2 in the central garden in the late season (increase), N1 in the southern garden in the early season (decrease), and C1 in the southern garden in the early season (decrease). None of the 10 combinations with an escape or avoidance pattern had credible intervals that excluded zero in both seasons. The magnitude of change also differed among gardens: Δ*g*_sw_ (mol m^−2^ s^−1^) ranged from −0.019 to 0.009 in the northern garden, from −0.019 to 0.024 in the central garden, and from −0.032 to 0.021 in the southern garden.

### Narrow-sense heritability across populations and gardens

Narrow-sense heritability was low overall: posterior median ℎ^2^ranged from 0.001 to 0.026 in the early season and from 0.004 to 0.087 in the late season (Fig. 5). ℎ^2^was higher in the late than in the early season in 13 of the 14 population-by-garden combinations (the 95% credible interval of the difference excluded zero), and this pattern was more pronounced in the central and southern gardens (median late minus early ℎ^2^ averaged across populations: 0.015 in the northern garden, 0.026 in the central garden, and 0.044 in the southern garden). For example, in the southern garden, late season ℎ^2^ was highest for N1 (0.087), C2 (0.086), and N2 (0.061), whereas early season ℎ^2^ was below 0.03 for all populations. Across all three gardens, population C1 had ℎ^2^ near zero in both seasons (never above 0.007).

### The relationship between Δ*g*_sw_ and ℎ^2^

We examined the relationship between Δ*g*_sw_ and ℎ^2^ using linear regression. For each posterior draw, there were 14 observations within each season. The posterior slopes (regression of Δ*g*_sw_, in mol m^−2^ s^−1^, on ℎ^2^) were centered near zero in both seasons (Fig. 6). In the early season, the mean slope was −0.01 (95% CI −0.88 to 0.89) and the posterior probability that the slope was positive was 0.46. In the late season, the mean slope was 0.01 (95% CI −0.16 to 0.17) and the posterior probability that the slope was positive was 0.54. We therefore found no evidence for a positive or negative relationship between heritability and evolutionary change in either season. The credible intervals were especially wide in the early season, when ℎ^2^was uniformly low.

**Figure 6:**
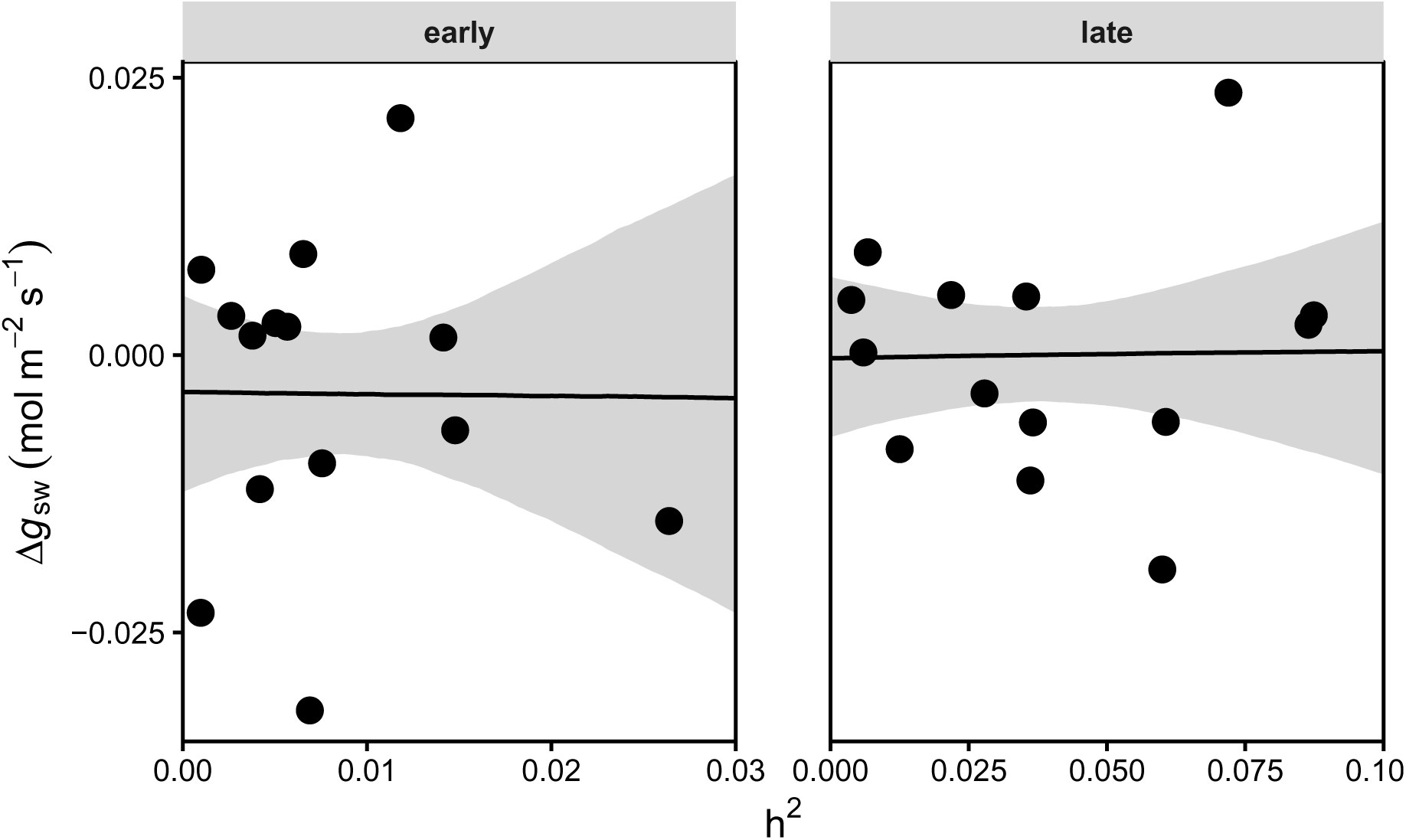
Relationship between heritability (ℎ^2^) and evolutionary change in Δ*g*_sw_ (mol m^−2^ s^−1^) across early and late seasons. Black lines represent linear regressions between ℎ^2^ and Δ*g*_sw_, and shaded areas indicate the 95% credible interval of the fitted relationships. Points are posterior medians for the 14 source-population-garden combinations.

## DISCUSSION

An interesting question in evolutionary ecology is whether rapid evolution can rescue populations from extreme climate events, and whether this potential differs across a species’ range (Gomulkiewicz and Holt, 1995; Bell and Gonzalez, 2009; Anstett et al., 2026). We used a resurrection experiment in three common gardens to ask whether stomatal conductance in *M. cardinalis* evolved in response to an extreme multiyear drought, and whether any change matched the predictions of drought escape (higher early season and lower late season *g*_sw_) or drought avoidance (the opposite; Fig. 1). *g*_sw_ differed strongly among gardens, seasons, and years, but evolutionary change between cohorts was small: only 4 of 36 changes had 95% credible intervals that excluded zero, and no escape or avoidance pattern (6 and 4 of 18 population-by-garden combinations) was supported in both seasons. Heritability of *g*_sw_ was low, higher in the late season, and unrelated to the magnitude of evolutionary change. Below we discuss how the environment shapes variation in *g*_sw_, possible reasons for limited evolutionary change, and what the heritability estimates imply.

### Environment and plasticity shape variation in *g*_sw_

We used *g*_sw_ as a physiological trait to test for drought escape and avoidance, and we asked whether *g*_sw_ varied among seasons, gardens, and source populations. We found that the environment strongly affects *g*_sw_, with consistent differences among seasons and locations. Across most gardens in 3 collection years (2023–2025), *g*_sw_ in the early season was higher than late season values, suggesting greater potential for carbon gain early in the season, before seasonal drought begins (Kooyers, 2015; Abobatta, 2019). The effect of the garden environment is consistent with plastic responses reported in this species, in which plants in a hot, dry environment show higher early season *g*_sw_ than plants in a cooler, wetter environment (Sheth et al., 2026), and with the high plasticity of leaf-level gas exchange (Anstett et al., 2021; Branch et al., 2024b). An exception to this pattern was observed in the northern garden in 2025, where late season *g*_sw_ was higher than early season values. This reversal may reflect unmeasured environmental variation, such as short-term weather changes, microclimatic differences, or shifts in the surviving plant composition due to differential mortality among populations and families in 2025. Differences in plant developmental stages at the time of measurement could also affect the data (Sheth et al., 2026) and legacy effects of climate from previous years on the phenotypes of perennial plants (Branch et al., 2024a).

Source populations differed significantly in *g*_sw_ when grown in the same gardens. For example, in the southern garden, the southern populations (S1, S2) exhibited significantly lower *g*_sw_ than the northern populations (N1, N2) in the early season of 2023 (difference in mean *g*_sw_: −0.06 (95% CI −0.08 to −0.05) mol m^−2^ s^−1^) and 2024 (−0.15 (95% CI −0.18 to −0.12) mol m^−2^ s^−1^), but not in the late season or in 2025. This may reflect the hotter, drier conditions in the southern garden. This pattern is consistent with local adaptation via drought avoidance, whereby populations from warmer, drier southern source regions have evolved more conservative stomatal behavior to limit water loss, while northern populations lack this selective history. Physiological traits of *M. cardinalis* vary clinally with latitude (Muir and Angert, 2017), and populations from different regions enter drought with different baseline leaf anatomies and drought strategies (Anstett et al., 2021; Branch et al., 2024b). Adaptive differences among populations may therefore be context dependent. Significant pairwise contrasts among populations were more frequent in the early season (49.2%) than in the late season (23.7%). Population-level differences in *g*_sw_ were therefore more pronounced in the early season than in the late season. We also found that *g*_sw_ differed much less among populations than other traits did in this experiment, such as leaf dry matter content, flowering time, and size at first flower (Sheth et al., 2026). In field gardens, some traits show less genetic differentiation among populations than the same traits in controlled conditions, likely because of greater environmental variation (Angert et al., 2011; Sheth and Angert, 2016; Muir and Angert, 2017; Nelson et al., 2021; Branch et al., 2024b). Genotype-by-environment interactions are important because the phenotypic effects of alleles in the field can be difficult to predict from studies in controlled conditions (El-Soda et al., 2014). Together, these points highlight the importance of environmental conditions.

Across the three collection years, early season *g*_sw_ declined in the northern and southern gardens. Temperature and soil moisture data from the gardens indicate that conditions remained consistently hot and dry during the study period (Fig. 2), suggesting that this decline may reflect cumulative physiological stress under persistently hot conditions; however, we cannot rule out other causes, such as ontogenetic changes, because plants also increased in size across years. Long-term studies in multiple common gardens show that warming and drying can erode the performance of locally adapted genotypes of plant species (Anderson et al., 2025). In contrast, the central garden showed a slight, non-significant increase in early season *g*_sw_ from 2024 to 2025. This may reflect a climate in the central garden that is more favorable for *M. cardinalis*. These results collectively highlight the importance of the local environment. The strong sensitivity of *g*_sw_ to local conditions is critical for interpreting the absence of clear evolutionary change, which we address in the next section.

### Little evidence for evolution of drought escape or avoidance

Plants can respond to drought with different strategies, such as drought escape or drought avoidance (Kooyers, 2015; Abobatta, 2019; Branch et al., 2024a; Branch et al., 2024b; Sheth et al., 2026). We asked whether *g*_sw_ evolved between ancestral and descendant cohorts in the direction predicted by either strategy. We hypothesized that drought escape would be associated with higher *g*_sw_ early in the season and lower *g*_sw_ late in the season, whereas drought avoidance is expected to show the opposite pattern, with lower *g*_sw_ early and higher *g*_sw_ late in the season (Franks et al., 2007, 2014; Hamann et al., 2018). We found no clear evidence for evolutionary change in *g*_sw_ in *M. cardinalis*. Although 6 of the 18 population-by-garden combinations showed patterns consistent with the evolution of increased escape (early Δ*g*_sw_ > 0 and late Δ*g*_sw_ < 0) and 4 with avoidance (opposite trend), none had 95% credible intervals that excluded zero in both seasons. The data thus do not support our hypothesis (Abobatta, 2019): *M. cardinalis* did not show clear evolutionary change in *g*_sw_ consistent with either drought response strategy between 2010 and 2017. This contrasts with a previous study on this species, which found that lower specific leaf area and later flowering evolved in central and southern regions, with weak or no change in the northern range, consistent with a shift from escape toward avoidance (Anstett et al., 2021). This difference may reflect that the earlier experiment grew plants in a greenhouse under controlled conditions (well-watered and terminal drought treatments). The authors suggested that plasticity plays an important role because anatomical or biochemical changes could reduce water loss without altering stomatal conductance (Anstett et al., 2021). Another study found that neither northern nor southern populations changed in photosynthesis or stomatal conductance under dry conditions after the drought, despite clear anatomical changes, and that stomatal conductance of northern populations changed clearly under wet conditions (Branch et al., 2024b). These studies show that *g*_sw_ responses depend on environmental and experimental conditions rather than being uniform. Escape and avoidance traits can evolve together (Kooyers, 2015), so a combination of responses can weaken the signal in any single trait. Finally, studies show that climate from earlier years can explain the direction of trait evolution better than conditions in the collection year, so conflicting selection among years and climatic variability can affect the detection of evolutionary change (Mulder et al., 2017; Branch et al., 2024a).

### Possible reasons for limited evolutionary change

Two main reasons may explain the absence of clear evolutionary change in *g*_sw_. First, the time interval between these cohorts may have been too short to capture strong evolutionary change. Evolution may need more time to occur, especially if this trait is governed by a complex, highly pleiotropic genetic architecture in which correlations among traits constrain adaptation (Etterson and Shaw, 2001; Gomulkiewicz and Houle, 2009). However, several previous resurrection studies, including some with *M. cardinalis*, show evidence of rapid evolutionary change in traits associated with drought adaptation (Franks et al., 2007; Franks and Weis, 2008; Franks, 2011; Hamann et al., 2018; Dickman et al., 2019; Anstett et al., 2021; Kooyers et al., 2021; Branch et al., 2024b), so time alone may not explain our result; other factors, such as constraints on *g*_sw_, may also be involved.

Second, *g*_sw_ may exhibit high phenotypic plasticity, allowing *M. cardinalis* to respond rapidly to environmental fluctuations. Because such plasticity can help buffer fitness against environmental change, it may reduce the strength of selection acting on this trait, thereby attenuating evolutionary change. Plasticity can be adaptive in variable environments, and its responses to drought have been documented in resurrection studies (Chevin et al., 2010; Franks, 2011; Anderson et al., 2012; Muir et al., 2025). However, plasticity is not uniformly adaptive across traits and regions, and can itself evolve (Anstett et al., 2021; Branch et al., 2024b). Our study shows high plasticity in *g*_sw_, with variation among gardens being larger than variation among populations within the same garden. This pattern suggests that local environmental conditions, rather than population-level genetic differences, were the primary driver of *g*_sw_ expression, and that variation in *g*_sw_ reflects plasticity more than genetic variance. Differences among populations may depend on the local garden environment. For example, population differences were comparatively small in the northern garden relative to the central and southern gardens. The differences among gardens may reflect variation in environmental sensitivity rather than consistent evolution among populations. The magnitude of differences between ancestral and descendant cohorts can like-wise vary across environments, and some differences are detectable only in particular environments (Karitter et al., 2026; Sheth et al., 2026). However, few studies have explored how adaptive plasticity and local adaptation may work together to rescue populations from severe drought (Chevin et al., 2010; Scheiner et al., 2017; Anderson and Wadgymar, 2020).

Evolutionary change in *g*_sw_ of *M. cardinalis* may therefore be limited by many factors or highly context-dependent. Local microclimatic conditions, such as soil moisture, temperature, and vapor pressure deficit, strongly influence the expression of *g*_sw_, and these conditions will continue to change. Understanding the physiological mechanisms that regulate *g*_sw_ in *M. cardinalis* would help clarify the timescale of evolutionary change in this trait, a promising direction for future research. Stomatal density (a developmental trait established during leaf formation) and stomatal aperture (a physiological trait regulated on shorter time scales via guard cell signaling) directly affect *g*_sw_, and changes in stomatal density can be compensated by changes in guard cell size and aperture (Shahinnia et al., 2016; Xie et al., 2022). Future research should therefore characterize variation in *g*_sw_ in response to different environmental factors and across multiple scales.

### Low, season-dependent heritability of *g*_sw_

Across populations and gardens, estimates of narrow-sense heritability (ℎ^2^) for *g*_sw_ were generally low, indicating that only a small proportion of phenotypic variation in *g*_sw_ is explained by additive genetic variance. This pattern is consistent with our finding of weak and inconsistent evolutionary change in Δ*g*_sw_: additive genetic variation appears to contribute little to evolutionary change in this trait. Low and imprecise heritability estimates under field conditions have been reported, particularly for physiological traits (Conner and Hartl, 2004; Hoffmann et al., 2017). In this study, ℎ^2^ was low, in contrast to the higher heritabilities reported for some traits and populations of *M. cardinalis* in controlled greenhouse settings (Sheth and Angert, 2016; Nelson et al., 2021), and similar contrasts have been reported in other systems (Geber and Griffen, 2003; Hoffmann et al., 2017). These low values likely reflect substantial environmental variance, non-additive genetic effects, or both, which reduce the proportion of variance attributable to additive genetic effects.

ℎ^2^ tended to be higher in the late season than in the early season, which suggests that the relative contribution of additive genetic variance increases under late season conditions. One explanation is that *g*_sw_ is more strongly influenced by environmental conditions in the early season, such as soil moisture, temperature, and competitive interactions during rapid growth. These factors can inflate environmental variance (*σ*^2^) and obscure genetic differences, leading to lower ℎ^2^ estimates. In contrast, late season conditions may be more stressful (drier conditions or more severe drought), reducing environmental variance and allowing additive genetic effects to be more clearly expressed. Classical evolutionary theory predicts that heritability is not fixed, but rather varies systematically between favorable and unfavorable conditions (Hoffmann and Merilä, 1999). This framework has been supported by multiple empirical studies (Bemmels and Anderson, 2019; Karitter et al., 2026). High phenotypic plasticity across small-scale environmental gradients may obscure the expression of genetic differences. For example, although genetic variation in phenology from experiments significantly predicted wild phenology across Eurasia, local variation in collection dates within a region was not explained by genetic variance in experiments (DeLeo et al., 2026). This suggests that environmental context, including small-scale or short-term environmental variation, can play a key role in the apparent expression of heritability. We observed that populations differed in heritability: mean late season ℎ^2^across gardens was highest for N1 (0.074) and C2 (0.060) and lowest for C1 (0.005). This pattern may reflect differences among populations in genetic variation, evolutionary history, and other genetic characteristics. Field data estimates of ℎ^2^ are known to be sensitive to environmental heterogeneity, which can inflate environmental variance and reduce additive genetic effects (Conner and Hartl, 2004; Hoffmann et al., 2017). Strong environmental influences, rather than a lack of genetic variation, may therefore explain the low and imprecise heritability estimates in this study.

### Heritability did not predict evolutionary change

We found no evidence that heritability affects evolutionary change across populations: the posterior slopes of Δ*g*_sw_ on ℎ^2^ were centered near zero in both seasons, and their credible intervals were wide (Fig. 6). Several factors may explain this lack of relationship. First, evolutionary change may be shaped by variation in past selection rather than current genetic variance. In our study, Δ*g*_sw_ reflects the cumulative outcome of selection acting between 2010 and 2017, whereas our estimates of ℎ^2^ are derived from recent environmental conditions (2023 to 2025). If the strength or direction of selection varied across this period, current heritability estimates may not accurately represent the genetic variance available at the time when selection occurred. Second, heritability depends strongly on the environment, which can affect the relationship between ℎ^2^ and Δ*g*_sw_, particularly if different gardens impose distinct environmental constraints on *g*_sw_ expression. According to the breeder’s equation, the response to selection is the product of heritability and the selection differential (*R* = ℎ^2^*S*) (Lande and Arnold, 1983; Falconer and Mackay, 1996; Sheth and Angert, 2016), so variable selection can further limit evolution even where additive variance exists (Grant and Grant, 1995; Sheth et al., 2026). These factors make it difficult to estimate quantitative parameters precisely under field conditions. Additive genetic variation alone may therefore not be sufficient to produce predictable evolutionary change in field environments.

## CONCLUSIONS

We used a seed resurrection approach in common gardens to examine evolutionary and plastic responses of stomatal conductance in *M. cardinalis* to drought. We evaluated the potential for adaptive responses to drought in the western United States between 2010 and 2017 and detected little evolutionary change in stomatal conductance. Moreover, we found no association between evolutionary change and heritability, so heritability did not explain differences in evolutionary change among populations and gardens. Instead, environmental context or historical selection could shape the dynamics of stomatal conductance, obscuring predictable responses to selection. These findings highlight a challenge in evolutionary ecology: even when genetic variation for a trait exists, its expression and evolutionary consequences may depend strongly on the environment. Future studies may require longer sampling periods and more detailed measurements of local climate conditions to determine whether long-term evolutionary responses emerge. In the future, we hope to use more data, traits, and variables to build predictive models of species’ responses to climate change and range shifts.

## ACKNOWLEDGMENTS

We thank Andy Alvarez, Kayla Bain, Gabe Bradshaw, Bryn Callie, Karina Cerda, Joanna Clines, Sulma Correa, Aeran Coughlin, Brooke Crose, Mia Cunningham, Maya Dingle, Lydia Duran, Anna Easton, Eve Eddy, Giovanni Gil, Mia Hanson, Ayleen Herrera, Allie Hoff, Nathanael Larson, Catherine Laufenberg, Yazmin Lommel, Eduardo Mendez, Camille Movafagian, Rachel Muir, Tyana Ortiz, Cristina Payst, Yoanna Rendon, Tadd Sexton, Emily Shank, Justin Shorty, Scarlet Steel, Everett Strain, Diana Tataru, Austyn Tavernier, Anika Vaidya, Emma Vtipilthorpe, Mia Wiegmann, Vincent Williams, Emma Wilson, Rachel Wooliver for help with data collection. We are grateful to greenhouse staff at the University of Oregon, San Diego State University, and the University of California, Merced, for support with plant growth facilities. David Lipson, James Bourdon, Alexander Powell, Jacob Keeton, and Pablo Bryant at the Santa Margarita Ecological Reserve; Mieko Aoki and Hal Cushman at the Friends of Buford Park and Mt. Pisgah Native Plant Nursery; and Joanna Clines from the Sierra National Forest provided permission and tremendous logistical and infrastructural support for this project. We thank Bonnie Bladen and Ray Laclergue for advice on garden design and pest management. Eddie Barragán and Freyja Moser provided feedback on this manuscript.

This work was supported by National Science Foundation projects DEB-2131815 (S.N.S.), 2131816 (J.D.), 2131817 (C.D.M.), 2131818 (J.P.S.), and 2131819 (L.F.-R.); a Research Capacity Fund (HATCH) project award (7002993) from the US Department of Agriculture’s National Institute of Food and Agriculture (S.N.S.); the Office of the Vice Chancellor for Research and Graduate Education at the University of Wisconsin-Madison with funding from the Wisconsin Alumni Research Foundation; North Carolina State University Goodnight Early Career Innovators Award (S.N.S.). Part of this research was supported by the Sierra Nevada Research Station–Yosemite Field Station (https://doi.org/10.21973/N3V36C).

## AUTHOR CONTRIBUTIONS

S.N.S., C.D.M., J.D., L.F.-R., and J.P.S. conceptualized and designed the study and obtained funding to support the work. D.W. helped develop protocols for data collection. All authors helped collect the data. C.D.M. and D.W. analyzed the data and wrote the original draft of the manuscript. All authors reviewed and approved the manuscript.

## DATA AVAILABILITY STATEMENT

Custom scripts and data are available on a GitHub repository (https://github.com/cdmuir/persist-gsw).

## SUPPORTING INFORMATION

Additional supporting information may be found online in the Supporting Information section at the end of the article.

- Appendix S1: Supplemental tables and figures

## APPENDIX S1

**Table S1:** Sampling time (first and last date of \gsw measurements) in each garden, season, and year. No early-season measurements were made in the central garden in 2023.

| Season | Year | Northern | Central | Southern |
|---|---|---|---|---|
| early | 2023 | 2023-05-23 to 05-31 |  | 2023-06-06 to 06-16 |
| late | 2023 | 2023-08-26 to 08-30 | 2023-08-22 to 08-24 | 2023-08-15 to 08-18 |
| early | 2024 | 2024-06-01 to 06-12 | 2024-05-23 to 05-27 | 2024-05-13 to 05-19 |
| late | 2024 | 2024-08-19 to 08-24 | 2024-08-12 to 08-17 | 2024-08-05 to 08-10 |
| early | 2025 | 2025-05-23 to 05-29 | 2025-05-14 to 05-20 | 2025-05-11 to 05-12 |
| late | 2025 | 2025-08-08 to 08-11 | 2025-08-04 to 08-05 | 2025-08-02 to 08-02 |

**Table S2:** *g*_sw_ estimates (mol m^−2^ s^−1^) by population, garden, year, and season. Values represent the posterior mean and 95% credible interval derived from model predictions. Early-season data were not collected for the central garden in 2023 due to logistical delays in transplanting, which resulted in insufficient plant growth for measurement at that time point. Negative lower bounds for southern garden 2025 credible intervals reflect high posterior uncertainty due to reduced sample size from increased mortality that year, rather than biologically meaningful negative conductance.

| Garden | Population | Year | Early-season *g*<sub>sw</sub> (95% CI) | Late-season *g*<sub>sw</sub> (95% CI) |
|---|---|---|---|---|
| Northern | N1 | 2023 | 0.351 (0.301-0.403) | 0.136 (0.075-0.196) |
| Northern | N1 | 2024 | 0.275 (0.228-0.322) | 0.102 (0.047-0.157) |
| Northern | N1 | 2025 | 0.092 (0.035-0.148) | 0.333 (0.273-0.395) |
| Northern | N2 | 2023 | 0.348 (0.300-0.398) | 0.155 (0.096-0.216) |
| Northern | N2 | 2024 | 0.289 (0.243-0.334) | 0.111 (0.057-0.165) |
| Northern | N2 | 2025 | 0.087 (0.030-0.141) | 0.319 (0.260-0.380) |
| Northern | C1 | 2023 | 0.420 (0.373-0.468) | 0.157 (0.099-0.216) |
| Northern | C1 | 2024 | 0.231 (0.186-0.274) | 0.100 (0.047-0.154) |
| Northern | C1 | 2025 | 0.085 (0.030-0.140) | 0.314 (0.256-0.373) |
| Northern | C2 | 2023 | 0.424 (0.375-0.474) | 0.158 (0.098-0.218) |
| Northern | C2 | 2024 | 0.246 (0.200-0.291) | 0.097 (0.043-0.152) |
| Northern | C2 | 2025 | 0.089 (0.032-0.144) | 0.328 (0.270-0.389) |
| Northern | S1 | 2023 | 0.436 (0.388-0.485) | 0.142 (0.084-0.202) |
| Northern | S1 | 2024 | 0.228 (0.184-0.272) | 0.099 (0.046-0.152) |
| Northern | S1 | 2025 | 0.072 (0.017-0.126) | 0.306 (0.250-0.365) |
| Northern | S2 | 2023 | 0.398 (0.350-0.447) | 0.148 (0.090-0.207) |
| Northern | S2 | 2024 | 0.268 (0.223-0.313) | 0.091 (0.038-0.144) |
| Northern | S2 | 2025 | 0.090 (0.035-0.144) | 0.302 (0.245-0.362) |
| Central | N1 | 2023 |  | 0.407 (0.250-0.537) |
| Central | N1 | 2024 | 0.218 (0.089-0.329) | 0.107 (0.027-0.199) |
| Central | N1 | 2025 | 0.344 (0.222-0.473) | 0.255 (0.100-0.440) |
| Central | N2 | 2023 |  | 0.427 (0.279-0.557) |
| Central | N2 | 2024 | 0.240 (0.115-0.352) | 0.115 (0.034-0.203) |
| Central | N2 | 2025 | 0.332 (0.207-0.467) | 0.251 (0.096-0.433) |
| Central | C1 | 2023 |  | 0.471 (0.321-0.606) |
| Central | C1 | 2024 | 0.240 (0.121-0.350) | 0.107 (0.031-0.190) |
| Central | C1 | 2025 | 0.392 (0.269-0.520) | 0.219 (0.070-0.402) |
| Central | C2 | 2023 |  | 0.444 (0.298-0.576) |
| Central | C2 | 2024 | 0.258 (0.139-0.372) | 0.102 (0.018-0.190) |
| Central | C2 | 2025 | 0.412 (0.286-0.541) | 0.207 (0.057-0.389) |
| Central | S1 | 2023 |  | 0.494 (0.350-0.627) |
| Central | S1 | 2024 | 0.231 (0.115-0.345) | 0.095 (0.020-0.178) |
| Central | S1 | 2025 | 0.354 (0.230-0.485) | 0.219 (0.065-0.398) |
| Central | S2 | 2023 |  | 0.510 (0.360-0.645) |
| Central | S2 | 2024 | 0.239 (0.124-0.352) | 0.111 (0.031-0.197) |
| Central | S2 | 2025 | 0.369 (0.244-0.500) | 0.232 (0.082-0.415) |
| Southern | N1 | 2023 | 0.685 (0.644-0.726) | 0.097 (0.048-0.144) |
| Southern | N1 | 2024 | 0.502 (0.436-0.568) | 0.209 (0.159-0.259) |
| Southern | N1 | 2025 | 0.044 (-0.158-0.244) | 0.046 (-0.101-0.197) |
| Southern | N2 | 2023 | 0.694 (0.654-0.735) | 0.105 (0.056-0.153) |
| Southern | N2 | 2024 | 0.510 (0.446-0.573) | 0.206 (0.158-0.254) |
| Southern | N2 | 2025 | 0.113 (-0.032-0.258) | 0.051 (-0.059-0.159) |
| Southern | C1 | 2023 | 0.636 (0.597-0.675) | 0.096 (0.047-0.143) |
| Southern | C1 | 2024 | 0.429 (0.377-0.481) | 0.260 (0.218-0.304) |
| Southern | C1 | 2025 | 0.104 (0.002-0.206) | 0.068 (-0.028-0.164) |
| Southern | C2 | 2023 | 0.620 (0.579-0.660) | 0.092 (0.042-0.141) |
| Southern | C2 | 2024 | 0.429 (0.374-0.484) | 0.246 (0.202-0.292) |
| Southern | C2 | 2025 | 0.080 (-0.043-0.204) | 0.049 (-0.056-0.154) |
| Southern | S1 | 2023 | 0.647 (0.609-0.687) | 0.099 (0.051-0.146) |
| Southern | S1 | 2024 | 0.376 (0.324-0.428) | 0.197 (0.155-0.241) |
| Southern | S1 | 2025 | 0.098 (0.007-0.185) | 0.075 (-0.015-0.164) |
| Southern | S2 | 2023 | 0.604 (0.566-0.643) | 0.091 (0.043-0.139) |
| Southern | S2 | 2024 | 0.344 (0.292-0.395) | 0.191 (0.149-0.234) |
| Southern | S2 | 2025 | 0.098 (0.008-0.187) | 0.076 (-0.014-0.165) |

**Figure S1:**
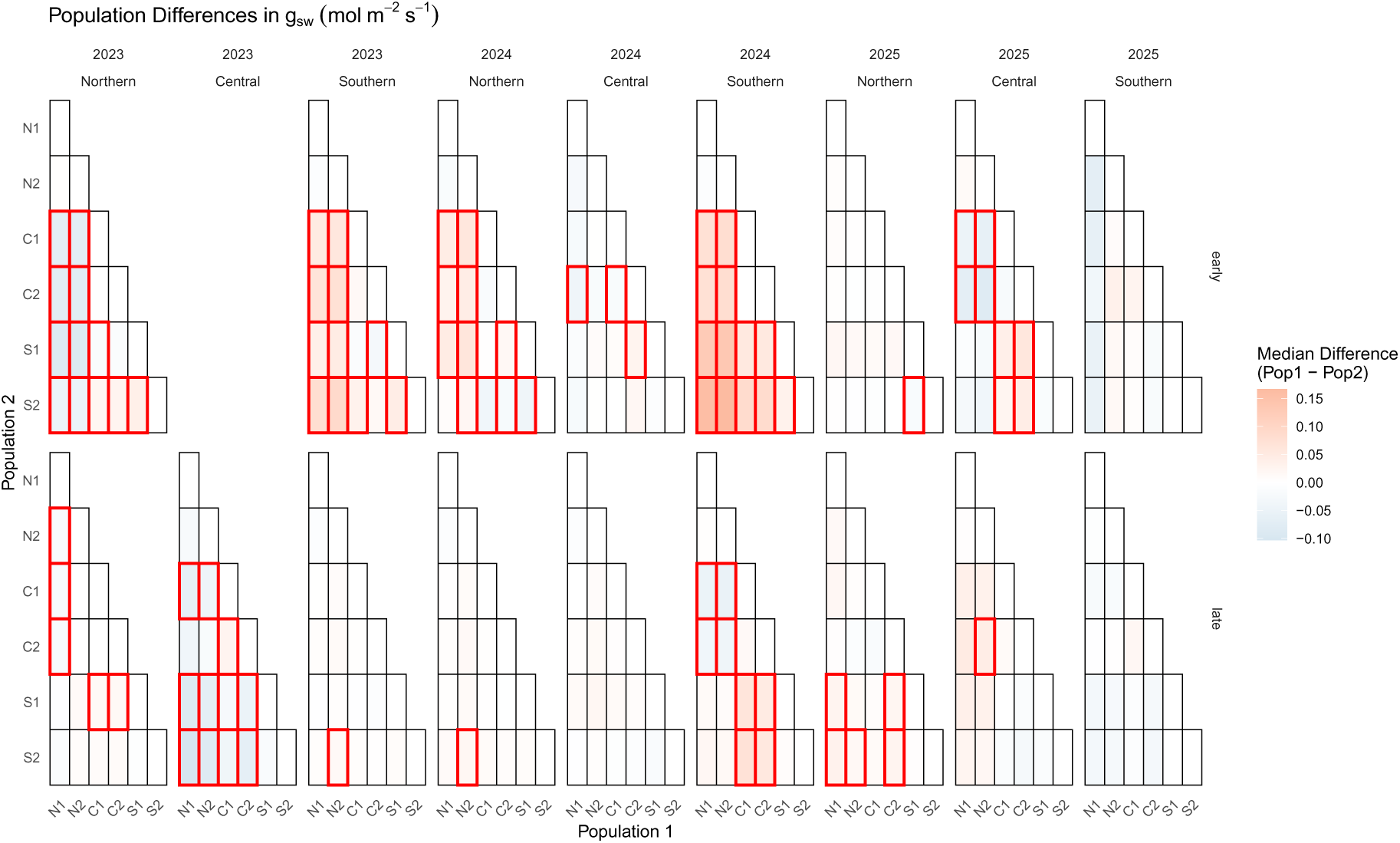
Differences in *g*_sw_ (mol m^−2^ s^−1^) among populations within each garden, season, and year. Red boxes indicate significant differences (the 95% credible interval of the difference excludes zero).

Pairwise comparisons of posterior differences in *g*_sw_ among populations within the same garden, season, and collection year revealed that median differences ranged between −0.10 and 0.17 mol m^−2^ s^−1^. Regarding statistical significance, the early season exhibited more significant pairwise contrasts than the late season, with 59 out of 120 comparisons (49.2%) being significant in the early season compared to 32 out of 135 (23.7%) in the late season. In 2023, only late season data are available for the central garden, and 11 of its 15 pairwise contrasts (73.3%) were significant. Across gardens, 38.9% of contrasts were significant in the northern garden, 30.7% in the central garden, and 36.7% in the southern garden. In 2025, the differences among populations in the southern garden were relatively small: 0 out of 30 contrasts were significant.

**Figure S2:**
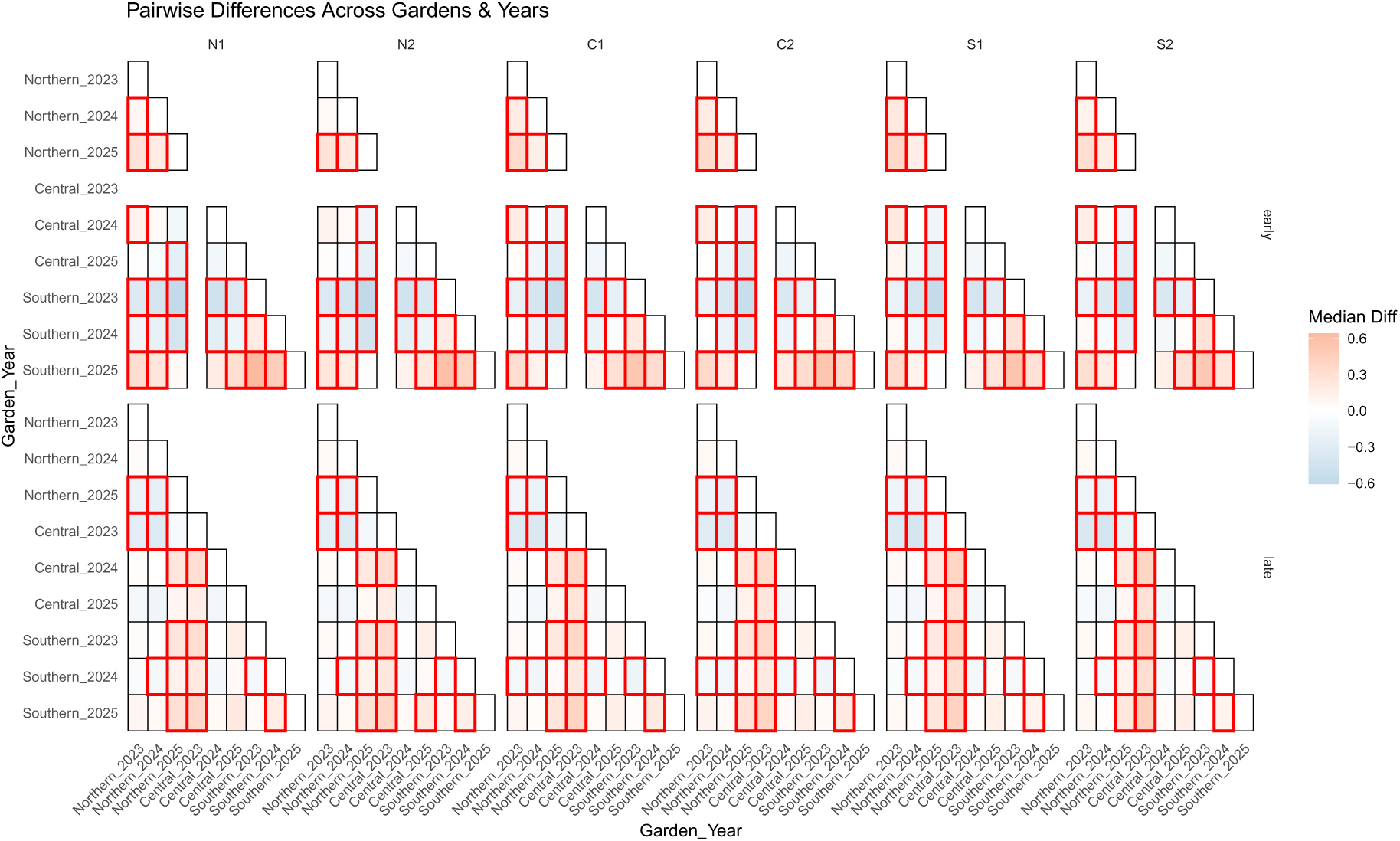
Differences in *g*_sw_ (mol m^−2^ s^−1^) between garden-year combinations within each population and season. Red boxes indicate significant differences (the 95% credible interval of the difference excludes zero).

This figure shows pairwise differences in *g*_sw_ between garden-year combinations within each population. Overall, more significant differences were observed in the early season (123 out of 168 contrasts, 73.2%) than in the late season (101 out of 216, 46.8%). The number of significant contrasts per population ranged from 19 (S2) to 22 (C2) out of 28 in the early season, and from 15 (N1) to 18 (C2 and S1) out of 36 in the late season. These patterns indicate that *g*_sw_ differed more among garden-year combinations in the early than in the late season.

